# Changes in bird migration phenology over six decades, a perspective from the Neotropical non-breeding grounds

**DOI:** 10.1101/2024.02.21.581429

**Authors:** Daniel A. Gutiérrez-Carrillo, Bryam Mateus-Aguilar, Camila Gómez, Carlos Daniel Cadena

## Abstract

Changes in the migration phenology of birds linked to global change are extensively documented. Longitudinal studies from temperate breeding grounds have mostly shown earlier arrivals in the spring and a variety of patterns during fall migration ^1,2^, yet no studies have addressed whether and how migration phenology has changed using data from the tropical non-breeding grounds. Understanding whether changes in migratory phenology are also evident in non-breeding sites is essential to determine the underlying causes of patterns documented in breeding areas. Using data from historical scientific collections and modern repositories of community-science records, we assessed changes in migration phenology of 12 Nearctic-Neotropical long-distance migratory birds in Colombia over six decades. We also explored whether shared breeding and non-breeding climatic niches explained variation in phenological patterns observed among species. All species showed shifts in spring (range −37 – 9 days from peak passage date) or fall (range −26 – 36 days) migration, but patterns differed among species in ways partly attributable to shared breeding or wintering climatic niches. Our results, although not yet broadly generalizable, suggest that birds use cues to time their migration at their non-breeding grounds which are most likely different to those they use on their breeding grounds. To better understand the effects of global change on biodiversity, exploring the underlying drivers of phenological changes with further research integrating more long-term datasets available through scientific collections and community science platforms should be a priority.

## Results and Discussion

We documented large phenological shifts in the spring (March – May) and fall (September – November) migration of birds over more than six decades in the Neotropics. By comparing thousands of historical and modern presence records, for a selection of 12 Nearctic-Neotropical long-distance migratory species which had at least 50 historical records in Colombia (Table 1), and for 99 species for which any historical data were available, we found that in 2009 – 2022, migrant birds arrived at their non-breeding grounds 7 days later in the fall (a delay of 1.1 days/decade) compared to 1908 – 1965 (Fig.1A). The observed mean phenological shifts increased when we included data for all 99 species with historical records (Suppl. Fig. 1). Observing such a shift based on data solely from the non-breeding period is remarkable given the view that cues observable only in the temperate zone (e.g. temperature and spring green-up^2^) are thought to largely drive the timing of migratory movements. Our findings thus open an unexplored avenue of research into the mechanisms at work in the tropics that drive phenological changes in Nearctic-breeding migratory birds.

**Figure 1.**
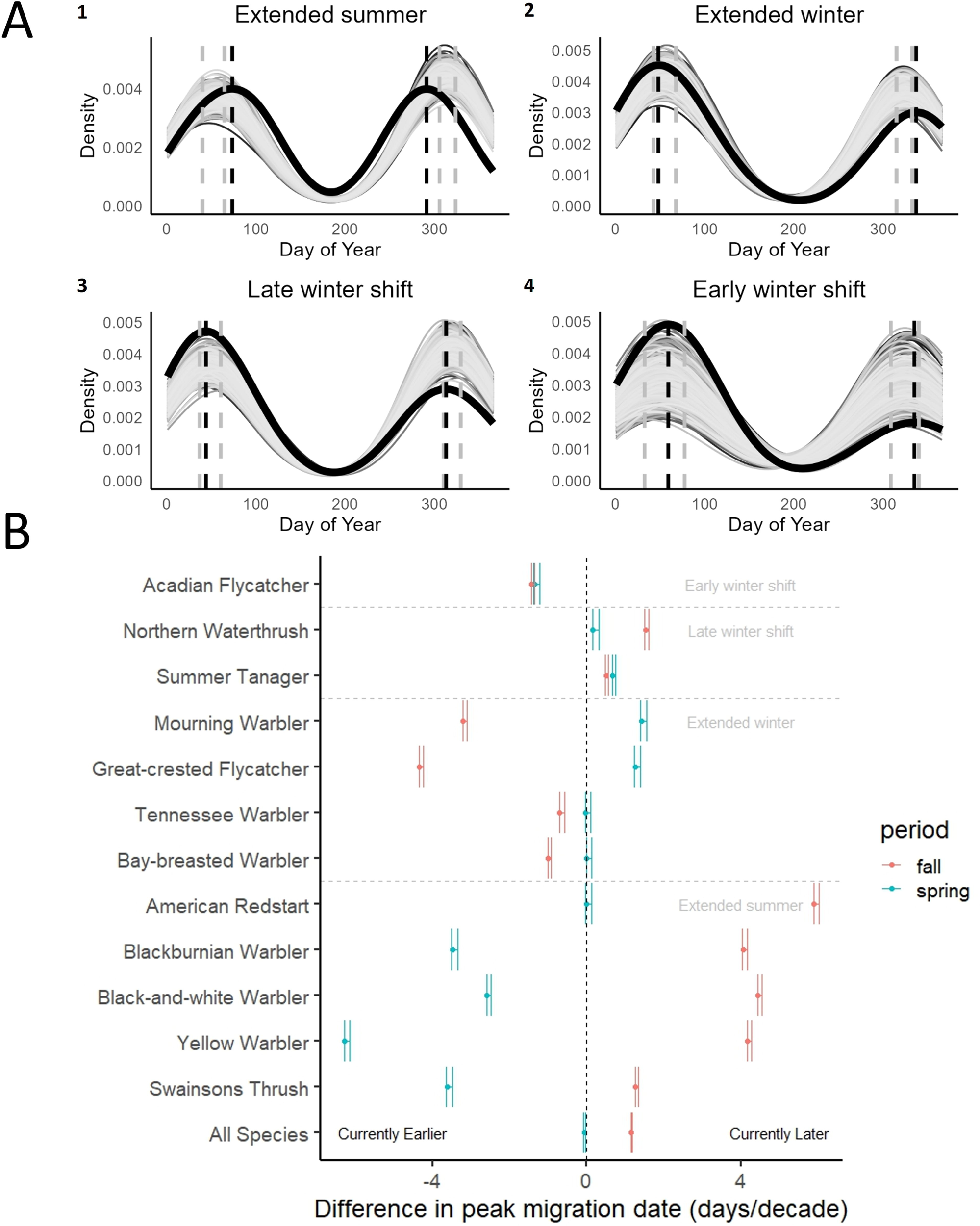
**A.** Phenological patterns comparing historical and modern data of 12 Nearctic-Neotropical long-distance migratory bird species detected in Colombia. The thick black curve shows the historical phenology, the light gray curves represent 1000 random draws from the modern dataset matching the historical number of records and reconstructing a distribution of modern density curves. Vertical dashed lines illustrate the peak migration dates for spring and fall (black dashed = historical) and the minimum and maximum estimates of peak dates from the modern random draws of data (gray dashed = modern). Pannels 1-4 illustrate the four phenological categories detected in this study: extended summer (i.e., species currently arrive to the non-breeding grounds later in the fall and leave them earlier in the spring), extended winter (i.e., species currently arrive to the non-breeding grounds earlier in the fall and leave them later in the spring), late winter shift (i.e., the whole non-breeding period shifted to a later date), and early winter shift (i.e. the whole non-breeding period shifted to an earlier date). **B.** Phenological shifts and confidence intervals estimated for 12 individual species. Difference in peak migration dates and their confidence intervals were estimated as the range of Δ = modern – historical for each random iteration and are presented as days/decade. Thus, negative values indicate current earlier dates and positive values indicate current later dates. Related to Figure S1 and S2.

**Table 1.** Number of historical and modern records available for each species and their classification of phenological shift category.

When analyzing each species separately, we found that the magnitude of the overall pattern observed with all records was not shared by all species (Figs.1A-B, Suppl. Fig. 2). Blackburnian Warbler (*Setophaga fusca*), Black-and-white Warbler (*Mniotilta varia*), Yellow Warbler (*S. petechia*), American Redstart (*S. ruticilla*), and Swainson’s Thrush (*Catharus ustulatus*), shared an ‘extended summer’ pattern, with individuals leaving their breeding grounds later in the fall and then returning earlier in the spring, and these species showed the largest magnitudes of change (Δ = modern - historical; mean Δ spring = −19 days or −3.1 days/decade, range −37 – 0 days in six decades; mean Δ fall = 24 days or 4 days/decade, range 7 – 36 days in six decades, Figs. 1A-B). However, Bay-breasted Warbler (*S. castanea*), Tennessee Warbler (*Leiothlypis peregina*), Great-crested Flycatcher (*Myiarchus crinitus*), and Mourning Warbler (*Geothlypis philadelphia*) showed an ‘extended winter’ pattern, where birds arrived earlier in the fall and left later in the spring (mean Δ spring = 4 days or 0.6 days/decade, range 0 - 8 days in six decades; mean Δ fall = −13 days or −2.1 days/decade, range −25 – −4 days in six decades, Figs.1A-B). Summer Tanager (*Piranga rubra*) and Northern Waterthrush (*Parkesia noveboracensis*) shared a pattern in which their entire winter period shifted to a later date, arriving later in the fall and leaving later in the spring (mean Δ spring = 3 days or 0.5 days/decade, range 1 - 4 days in six decades; mean Δ fall = 6 days or 1 day/decade, range 3 – 9 days in six decades, Figs. 1A-B). Lastly, Acadian Flycatcher (*Empidonax virescens*) showed a unique pattern of shift in its entire non-breeding period, arriving 8 days earlier in the fall (1.3 days/decade), and leaving 8 days (1.3 days/decade) earlier in the spring (Figs. 1A-B).

Our results, uniquely based on data from the Neotropical non-breeding grounds, are consistent with patterns revealed by studies evaluating phenological shifts in the temperate zone which show evidence of earlier arrivals to the breeding grounds during spring and varying trends for fall departures ^1,3–12^. Aside from a study including data from the Southern Hemisphere (mainly Australasia) and which highlighted a lack of information from the Neotropics ^7^, to our knowledge there has been no other research exploring temporal shifts in bird migration phenology from the Neotropical non-breeding period. Recent studies from North America report spring advancements of similar magnitudes to those we observed: between 0.5 and 2.9 days per decade for 9 species over 60 years (1961-2018) ^1^, and a spring advancement of 2.2-2.4 days per decade at a global scale and 0.4 to 1.0 days in North America over 38 years (1978-2016) ^8^. In terms of individual species, our results agree with those of Horton et al. (2023) for American Redstart and Yellow Warbler, which were both shown to be arriving on their breeding grounds ∼1 day/decade earlier in the spring and generally later in the fall, though confidence intervals overlapped with 0 ^1^. Tennessee Warbler, Northern Waterthrush, Mouring Warbler, and Bay-breasted Warbler have all been shown to be arriving on the breeding grounds ∼1 day/decade earlier in the spring while we detected later spring departures from the non-breeding grounds for these species (Fig.1B). Although later departures from the non-breeding grounds can result in faster overall migrations if birds are in good-quality habitats^13^, more study is needed to confirm whether this is indeed the case for the aforementioned species. What we can ascertain is that patterns of global change are heterogeneous and thus responses and drivers of change in temperate regions are not necessarily the same as those acting at lower latitudes^4,7^. It is therefore both surprising and enlightening that we find similarities with studies from the temperate zone. Our work is also consistent with prior temperate-zone studies revealing that phenological shifts in migration vary among species ^3,5,6^, which raises questions about the mechanisms at work within the Neotropics that may drive such changes in different species and populations ^7,14^. Having said this, we acknowledge that the12 species with information in this study represent a low sample size to provide broad generalizable conclusions, so we urge future research involving other species to test and build on our findings.

We hypothesized that species with equivalent patterns of phenological shift might be responding similarly to common environmental cues for breeding and migration^15^ owing to them potentially sharing breeding or non-breeding climatic niches. If this were the case, then we would expect that species sharing phenological shift patterns would also show a high degree of overlap in their breeding or non-breeding climatic niche compared to the niche overlap between species with different phenological patterns. We characterized niches based on seven uncorrelated climatic variables used in earlier work to describe Neotropical migratory bird climatic niches ^15^: elevation, aspect, slope, maximum temperature, minimum temperature, precipitation seasonality, and temperature seasonality.

Species sharing the extended summer pattern had higher mean overlap in their breeding climatic niche compared to their winter climatic niche (Fig.2A, mean D_summer_ = 0.61 ± 0.12 SD, mean D_winter_ = 0.49 ± 0.22 SD, P = 0.09, n = 10 comparisons, 5 spp, see also Suppl. Figs.3-4). Conversely, species sharing the extended winter pattern had a higher winter niche overlap compared to their breeding climatic niche overlap (mean D_winter_ = 0.61 ± 0.19 SD, mean D_summer_ = 0.44 ± 0.27 SD, P = 0.01, n = 6 comparisons, 4 spp see also Suppl. Figs.3-4). The two species sharing the late winter shift pattern had a low breeding niche overlap value of D_summer_ = 0.04, and a high winter niche overlap of D_winter_ = 0.72 (Fig. 2A, Table 1, see also Suppl. Figs. 3-4). We did not detect large differences in climatic niche overlap between groups of species sharing phenological shift patterns (intra group) and those with different patterns (inter group), either in the breeding period (Fig. 2B, Suppl. Table 1, intra group mean D_summer_ = 0.52 ± 0.23 SD, inter group mean D_summer_ = 0.40 ± 0.27 SD, P = 0.94), or the non-breeding period (intra group mean D_winter_ = 0.55 ± 0.21 SD, inter group mean D_winter_ = 0.60 ± 0.18 SD, P = 0.18). Although our results apply only to the species we evaluated, the patterns we observed suggest that summer climatic niches may influence phenological shifts that imply a longer breeding period, potentially through an early onset of spring and later fall, whilst non-breeding climatic conditions may be influencing phenological shifts which imply a longer winter period, perhaps through shifts in the precipitation regime on the non-breeding grounds. More research is needed to further test these hypotheses and to assess the generality of our findings across more species.

**Figure 2.**
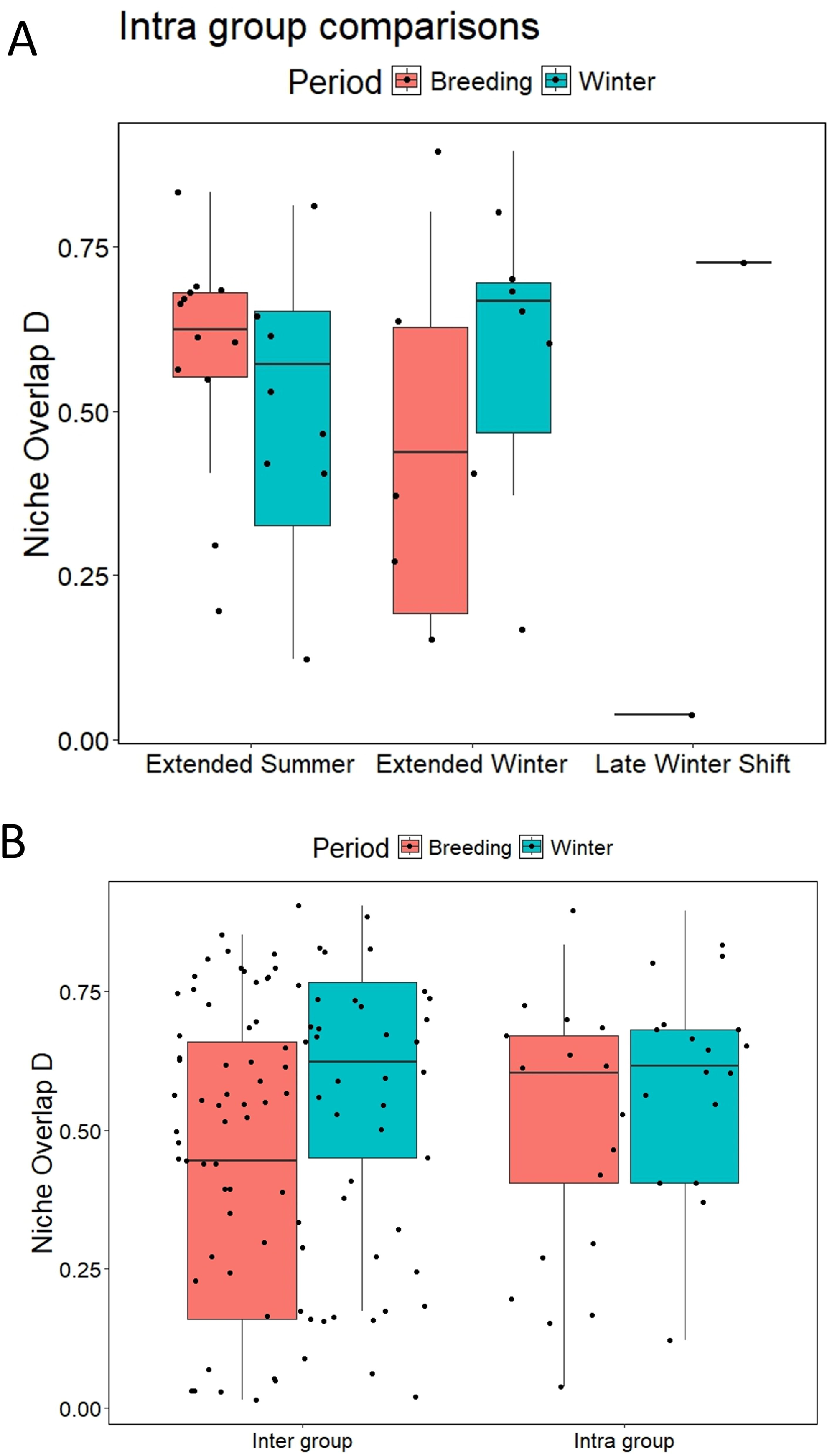
**A.** Climatic niche overlap of 12 Nearctic-Neotropical long-distance migratory bird species per phenological category. Species under the “extended summer” phenological category share a higher proportion of their breeding climatic niche (D_summer_ = 0.61 ± 0.12 SD) compared to their non-breeding niche (mean D_winter_ = 0.49 ± 0.22 SD), whilst species with an extended winter share a higher proportion of their non-breeding climatic niche (mean D_winter_ = 0.61 ± 0.19 SD) compared to their breeding niche (mean D_summer_ = 0.44 ± 0.27 SD). The two species sharing the late winter shift pattern show low breeding niche overlap (D_summer_ = 0.04), and high winter niche overlap (D_winter_ = 0.72). **B.** Comparisons of niche overlap between species sharing phenological shifts (intra-group) and those not sharing them (inter-group) did not show large differences in either the breeding or the non-breeding period. Related to Table S1 and Figures S3 and S4.

Both cue-driven environmental mechanisms (e.g. temperature, precipitation, photoperiod, vegetation greening) and response-driven organismal mechanisms (e.g. trophic level, migratory strategy, ecological specialization), which vary among species, may underlie phenological shifts ^2,16–18^. However, the cues to which migratory birds respond on their non-breeding grounds likely differ from those acting on the breeding grounds ^7^, particularly when these areas are distant. In North America, temperature appears to be the main cue-driven mechanism underlying shifts in spring arrivals and fall departures ^1,9,10^. Because seasonal temperature variation in northern South America is minimal, changes in temperature are unlikely drivers of the patterns we observed. Alternatively, changes in precipitation cycles may provide tangible cues to drive phenological shifts in migration to and from the Neotropical non-breeding grounds ^7,19^. Regular rain and drought cycles, as well as precipitation fluctuations associated with climatic phenomena such as El Niño and La Niña can affect resource availability in the tropics ^20–22^, and migratory birds respond to variation in precipitation on their non-breeding grounds by modifying their migration schedules ^23–26^.

Whether precipitation on the non-breeding grounds in turn relates to temperature or other environmental conditions on the breeding-grounds is not well understood, and how landbirds may shift their cue perception as their migration advances needs more study ^27,28^. We are unaware of research on organismal-driven cues (e.g. social foraging cues, individual breeding condition) potentially acting in the tropics and which may also be influencing phenological shifts in migration ^7^. Furthermore, the 12 species we analyzed in this study were those with enough historical data to allow meaningful comparisons across time, yet the patterns we uncovered are likely more general given that phenological shifts in migration have been detected in a broad range of Nearctic-Neotropical migratory species (Suppl. Fig. 1) ^1^. There is a broad and wide-open research avenue exploring how migratory species, populations and individuals respond to environmental and organismal cues throughout their annual cycle and whether these responses translate into adaptation to global change ^29^, or to increases in phenological mismatches ^30,31^.

Long-distance Nearctic-Neotropical migratory birds are responding to climate and global change by shifting their fall and spring migration schedules, both on the breeding ^1,8,10^ and non-breeding grounds (this study). Yet, the persistent and steep population declines of Nearctic Neotropical long-distance migrants ^32^ and the broad range of responses to global change in different species, suggest that neither plasticity or adaptability may be quick enough or uniform among species to respond to the rising pressures posed by a rapidly changing world ^2,8,30,31^. Phenological shifts in migration can accentuate mismatches between arrivals, departures, and resource peaks that migrants depend on during different periods of their annual cycle ^30,33–35^. The fitness and demographic consequences of such mismatches could drive declines in certain populations^34^.

Our study highlights the potential to address questions on a global scale by incorporating data from multiple sources. Biological collections provide a snapshot of biodiversity at specific times and locations and, thus, allow tracking of changes dating back decades or even centuries. However, historical sampling is not uniform across time or geographical locations in the global south. Multiple studies addressing changes in migration phenology in birds have been possible due to long-term monitoring through bird banding and standardized surveys. Such datasets are often limited in many tropical regions due to insufficient or poorly documented long-term biodiversity monitoring efforts. The growing popularity and usage of community science platforms have increased the accessibility to data that would otherwise have required extensive effort and resources to be obtained^36^. These data have proven sufficient to detect patterns over a wide diversity of timescales and geographic ranges^37^. Despite differences in the amount of data obtainable between biological collections and citizen science platforms, our results show how both sources of data can complement each other. We point out yet again the importance of a more complete understanding of the annual cycle of migratory organisms ^38,39^, where events occurring on the non-breeding grounds and Neotropical stopover sites are also affecting the phenological dynamics of migration.

## Acknowledgements

We thank Jose Vicente Rodríguez and Maria Isabel Moreno for sharing Conservation International’s historical dataset of bird records for Colombia (Ara Colombia). This study is part of the Colombia Resurvey Project which aimed to assess changes in bird abundance and diversity after a century of change since the American Museum of Natural History expeditions led by Frank Chapman in the early 1900’s. Dr. Nick Bayly and three anonymous reviewers provided helpful comments and discussion on earlier versions of the manuscript.

## Author contributions

CDC and CG designed the study. DAG, BMA and CG compiled and analyzed the data. All authors contributed to result interpretation and manuscript writing.

## STAR Methods

### Resource availability

#### Lead Contact

Camila Gómez

#### Material availability

https://github.com/camilgomo/Changes-in-bird-migration-phenology-over-six-decades

#### Data and code availability

https://github.com/camilgomo/Changes-in-bird-migration-phenology-over-six-decades

### Method details

#### Historical and modern data

We compiled all known historical records of Neotropical migratory birds in Colombia between 1908 and 1965 obtained from public data repositories of scientific collections (dataset available in GitHub: https://github.com/camilgomo/Changes-in-bird-migration-phenology-over-six-decades). The historical period, comprising close to six decades, was established as first, a period with enough data to make any sensible comparisons, and second, a period reflecting pre-industrialization and globalization environmental conditions, which exponentially increased after the 80’s (NOAA 2023, https://www.ncei.noaa.gov/access/monitoring/climate-at-a-glance/global/time-series/globe/land_ocean/ytd/12/1850-2023). We believe these human-induced changes have potentially exacerbated environmental and climatic variations experienced by migratory birds currently. Our complete historical database comprised information for 2455 specimens of 99 species from 31 scientific collections worldwide. For every record we verified species, date, locality, and geographic coordinates by accessing the original collection records ^40^. For the individual species comparisons, we selected species that had complete historical records for at least 50 individuals and removed austral migrants and species with both resident and migratory populations in Colombia due to potential ambiguity in their identification (e.g. *Vireo olivaceus, Tyrannus savana*). This reduced the historical dataset to 876 individuals of 12 species (dataset available in GitHub: https://github.com/camilgomo/Changes-in-bird-migration-phenology-over-six-decades). We used the global community science platform eBird ^41^ to obtain contemporary records of Neotropical migratory birds in Colombia between 2009 and 2022, for all Nearctic Neotropical migrants that winter or fly through Colombia, and for just the 12 species selected above in our historical dataset. After filtering for verified records, and eliminating duplicates, we gathered a total of 333,068 observations.

#### Exploration of potential biases in historical and modern datasets

We wanted to make sure that any patterns we observed in the migration phenology were not a result of temporal or spatial biases inherent to our datasets. We therefore explored whether there were temporal and spatial biases by plotting the availability of records by year, month and by latitude and longitude. Complementarily, we ran a Durbin-Watson test for autocorrelation between the number of records and latitude, longitude and year, using the R package *car*^42^. We did this both for the complete datasets, and for individual species. The historical dataset did not show any temporal bias or spatial biases, whereas the modern data showed a correlation with year (Suppl. Table 2, Suppl. Figs. 5-7). However, this is unsurprising since data recorded on the eBird platform has incremented annually since the platform was launched, so we continued with our analyses as described below.

#### Comparison of historical and modern migration phenology

We constructed density plots of migration phenology for the historical and modern datasets, both for all species together and for each species individually. We overlapped historical and modern phenology plots and determined the date of maximum density of migrants in Colombia during spring (peak abundance between March and May) and fall migration (peak abundance during September and November) by calculating the local density maximum for each period. We considered the day of maximum density to be a reliable metric of the general phenological pattern of a species ^43^. We then estimated the difference in peak dates by subtracting the modern from historical peak dates for fall and spring. Negative values mean fall or spring migration currently peaks earlier, and positive values mean fall or spring migration currently peaks later compared to 1908-1965 in Colombia.

Because we had such a broad difference in number of records between the historical and modern datasets (Table 1), which could potentially bias our comparisons, the modern phenology curves were generated by randomly drawing from the modern dataset the same number of historical records available for each species, then estimating the local maxima date and iterating this procedure 1000 times. The differences between the historical peak date of passage and each of the 1000 iterated modern peak dates, provided our confidence intervals for the phenological shifts observed during spring and fall migration. All analyses were carried out in R ^44^ and all code and data are available in GitHub (https://github.com/camilgomo/Changes-in-bird-migration-phenology-over-six-decades).

#### Breeding and non-breeding climatic niche overlap for species sharing phenological shift patterns

We used the R package ecospat ^45^ to estimate climatic niche overlap values for species which shared phenological patterns of migration shift in four categories based on the trends of the day of maximum density between historic and modern data: 1) extended summer (earlier spring departure and later fall arrival to the non-breeding grounds), 2) extended winter (later spring departure and earlier fall arrival to the non-breeding grounds), 3) late winter shift (the whole non-breeding period shifted to a later date) and 4) early winter shift (the whole winter period shifted to an earlier date).

We compiled monthly climatic variables for the breeding period (June – August) and the non-breeding period (December – February) using Worldclim data ^46^. Seven uncorrelated climatic variables were selected^15^, elevation, aspect, slope, maximum temperature, minimum temperature, precipitation seasonality, and temperature seasonality. Current breeding and non-breeding distributions for all species were obtained from eBird Status and Trends ^47,48^ by selecting the specific weeks that correspond to the breeding and non-breeding periods for our target species. We then filtered breeding and non-breeding ranges within the top 50% probability of occurrence of each species’ distribution range and then generated 5000 random points to be overlaid on the range and get presence xy coordinates for each species. Random breeding and non-breeding distribution coordinates were used to extract climate values for each species, and an additional 10000 random points were used to extract climatic background values for the whole of North America for the breeding period, and Central and South America for the non-breeding period).

Following Broennimann et al. ^45,49^ we conducted a multivariate comparison of climatic niche overlap between species which shared phenological shifts in historical and modern migration patterns. Climatic niche overlap is measured using Warren’s D statistic which varies from 0 (no climatic niche overlap), to 1 (climatic niches are exactly the same) ^15,49^. We compared values of niche overlap between species sharing the same phenological pattern (intra-group) and those not sharing them (inter-group). To determine whether mean D values were significantly different between phenological shift categories and both within and between category groups, we estimated the observed difference in mean values, and randomized each dataset to obtain a distribution of 999 random mean differences. The probability (P) associated with our observed difference in means was obtained by locating the observed difference within the random distribution of differences in means. All data and code are available form GitHub (https://github.com/camilgomo/Changes-in-bird-migration-phenology-over-six-decades).

### Key resources table

| RESOURCE | SOURCE | IDENTIFIER |
| --- | --- | --- |
| Deposited data |  |  |
| Raw historical records of Neotropical migratory birds recorded in Colombia between 1908 - 1965 | GBIF VertNet, Conservation International Colombia | https://www.gbif.org/<br>http://www.vertnet.org/ |
| Processed historical dataset of Neotropical migratory birds recorded in Colombia 1908 - 1965 | This study | https://github.com/camilgomo/Changes-in-bird-migration-phenology-over-six-decades |
| Raw modern observation records of Neotropical migratory birds in Colombia 2009-2022 | eBird | https://ebird.org/ |
| Processed modern dataset of Neotropical migratory birds in Colombia 2009-2022 | This study | https://github.com/camilgomo/Changes-in-bird-migration-phenology-over-six-decades |
| Breeding and non-breeding distribution rasters of neotropical migrants | eBird Status and trends | https://science.ebird.org/es/status-and-trends/download-data |
| Presence geographic coordinates for breeding and non-breeding climatic niche overlap | This study | https://github.com/camilgomo/Changes-in-bird-migration-phenology-over-six-decades |
| Monthly climatic variables | Worldclim | https://www.worldclim.org/ |
| Other |  |  |
| R Code and data for migration phenology comparisons | This study | GitHub |
| R Code and data for climatic niche overlap comparisons | This study | GitHub |

## Supplemental information

Document S1. Figures S1-S7 and Tables S1 and S2.

**Figure S1.** Phenological change graphs for ninety-nine migratory species recorded in Colombia between 1908 – 1965. The thick black curve shows the historical phenology, the light gray curves represent 1000 random draws from the modern dataset matching the historical number of records and reconstructing a distribution of modern density curves. Vertical black dashed lines indicate the peak dates of historical spring and fall passage, and the gray dashed lines show the minimum and maximum peak dates from the random draw of 1000 density curves from the modern dataset. Related to Figure 1.

**Figure S2.** Phenological change graphs for each of the 12 Nearctic-Neotropical long-distance migratory birds with more than 50 historical records assessed in this study. Extended summer: A) Black-and-white Warbler, B) Blackburnian Warbler, C) Yellow Warbler, D) American Redstart; E) Swainson’s Thrush. Extended Winter: F) Great Crested Flycatcher, G) Mourning Warbler, H) Bay-breasted Warbler, I) Tennessee Warbler. Late winter shift: J) Northern Waterthrush, K) Summer Tanager. Early winter shift: L) Acadian Flycatcher. The thick black curve shows the historical phenology, the light gray curves represent 1000 random draws from the modern dataset matching the historical number of records and reconstructing a distribution of modern density curves. Vertical black dashed lines indicate the historical peak dates of spring and fall passage, and the gray dashed lines show the minimum and maximum peak dates from the random draw of 1000 density curves from the modern dataset. Related to Figure 1.

**Figure S3.** Breeding climatic niche overlap between species that shared their phenological category. Extended Summer (A – J), Extended Winter (K – P), Late winter shift (Q). Niche overlap D values for each comparison shown in the box. Related to Figure 2 and Table S1.

**Figure S4.** Non-breeding climatic niche overlap between species that shared their phenological shift category. Extended Summer (A – J), Extended Winter (K – P), Late winter shift (Q). Niche overlap D values for each comparison shown in each box. Related to Figure 2 and Table S1.

**Figure S5.** Number of historical records available between 1908-1965, show that there is no detectable temporal bias in the availability of records. The high numbers between 1911 and 1914 correspond to the Natural History expeditions led by the American Museum of Natural History at that time (Chapman 1917). Related to STAR Methods.

**Figure S6.** Evaluation of the latitudinal and temporal distribution of historical (red) and modern (green) records of migratory species suggest there are no temporal and spatial biases in the data that could be influencing the patterns observed. Related to STAR Methods.

**Figure S7.** Maps of Colombia showing the geographic distribution of migratory bird records in the A. historical dataset, and B. modern dataset. Related to STAR Methods.

**Table S1.** Values of breeding climatic niche overlap (Dsummer), and B. non-breeding climatic niche overlap (Dwinter) estimated for all pairs of species in this study. Related to Figure 2 and Figure S3 and S4.

**Table S2.** Values of the Durbin-Watson autocorrelation test between the number of records of migratory birds and year, longitude and latitude, both for historical and modern datasets. Values around 2 of the D-W statistic show no autocorrelation of residuals. Values closer to 0 or to 4 show a positive or negative correlation respectively and are shown in bold. Related to STAR Methods and Figures S5 – S7.

