## Supplemental Tables & Figures for "Changes in bird migration phenology over six decades, a perspective from the Neotropical non-breeding grounds"

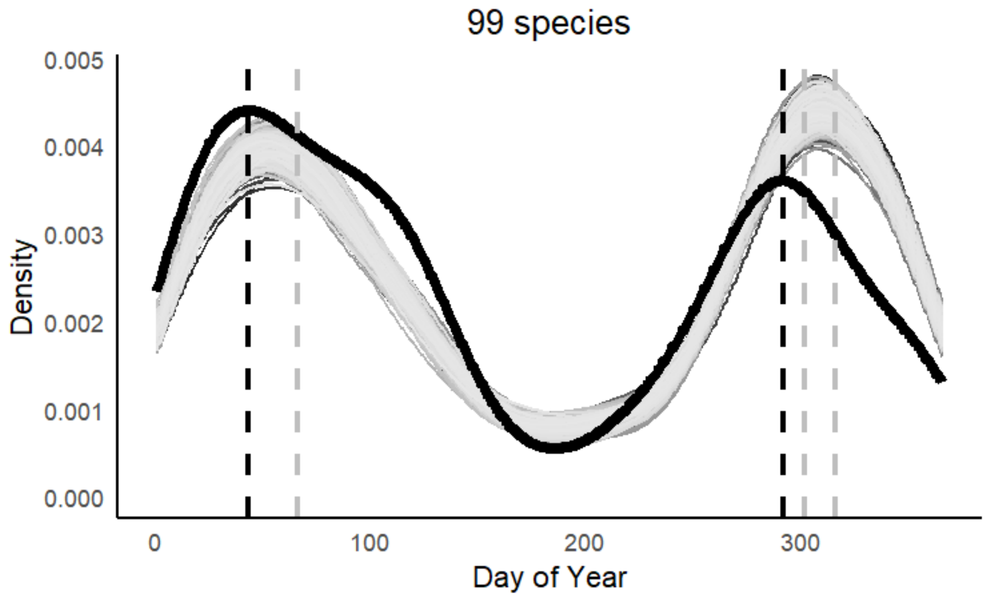


**Supplementary Figure 1.** Phenological change graphs for ninety-nine migratory species recorded in Colombia between 1908 – 1965. The thick black curve shows the historical phenology, the light gray curves represent 1000 random draws from the modern dataset matching the historical number of records and reconstructing a distribution of modern density curves. Vertical black dashed lines indicate the peak dates of historical spring and fall passage, and the gray dashed lines show the minimum and maximum peak dates from the random draw of 1000 density curves from the modern dataset. Related to Figure 1.

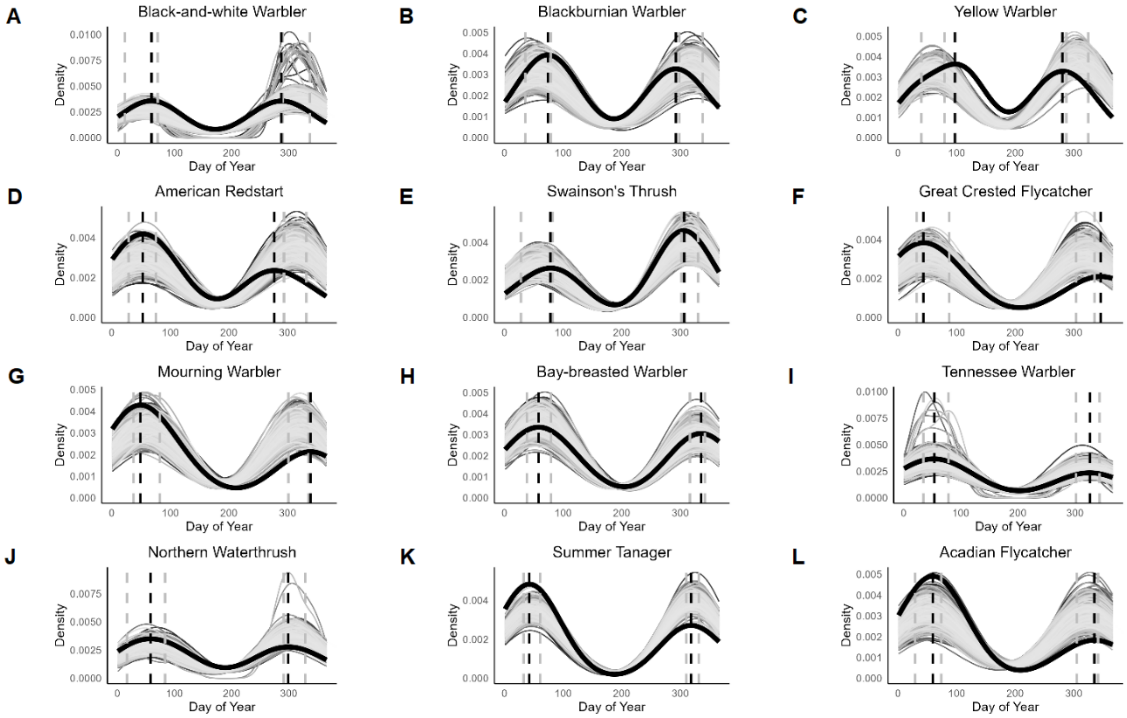


**Supplementary Figure 2.** Phenological change graphs for each of the 12 Nearctic-Neotropical long-distance migratory birds with more than 50 historical records assessed in this study. **Extended summer: A)** Black-and-white Warbler, **B)** Blackburnian Warbler, **C)** Yellow Warbler, **D)** American Redstart; **E)** Swainson's Thrush. **Extended Winter: F)** Great Crested Flycatcher, **G)** Mourning Warbler, **H)** Bay-breasted Warbler, **I)** Tennessee Warbler. **Late winter shift: J)** Northern Waterthrush, **K)** Summer Tanager. **Early winter shift: L)** Acadian Flycatcher. The thick black curve shows the historical phenology, the light gray curves represent 1000 random draws from the modern dataset matching the historical number of records and reconstructing a distribution of modern density curves. Vertical black dashed lines indicate the historical peak dates of spring and fall passage, and the gray dashed lines show the minimum and maximum peak dates from the random draw of 1000 density curves from the modern dataset. Related to Figure 1.

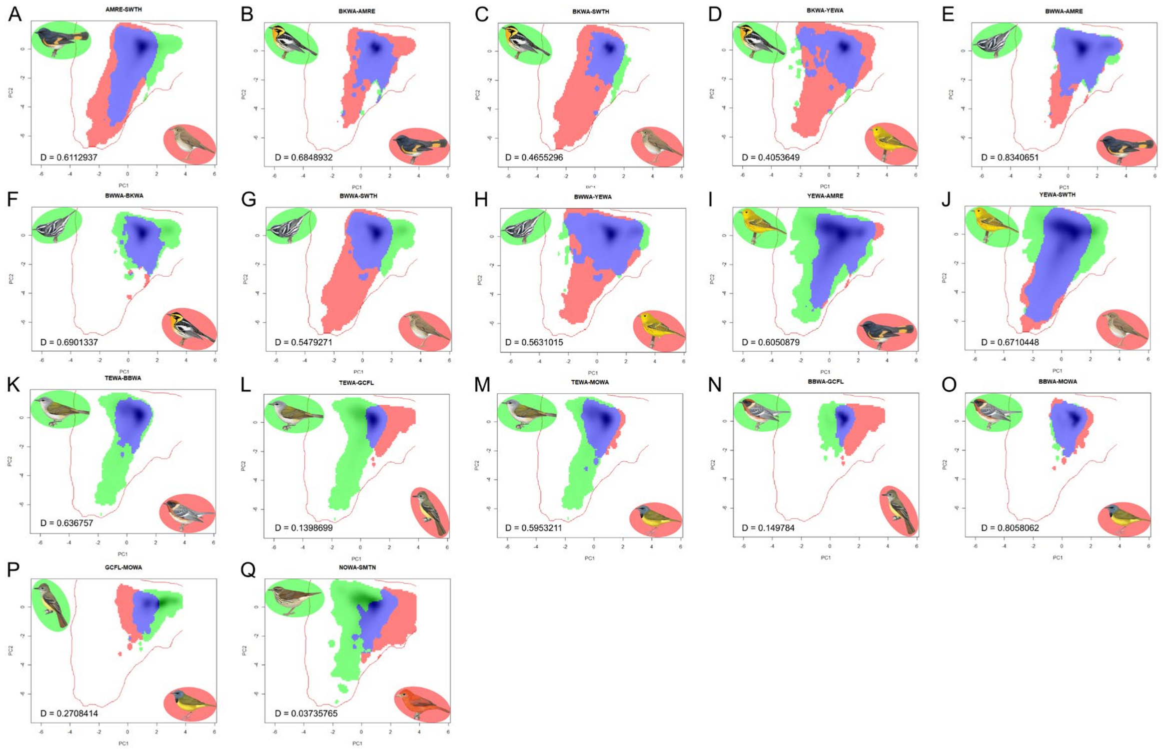


**Supplementary Figure 3.** Breeding climatic niche overlap between species that shared their phenological category. Extended Summer (A – J), Extended Winter (K – P), Late winter shift (Q). Niche overlap D values for each comparison shown in the box. Related to Figure 2 and Table S1.

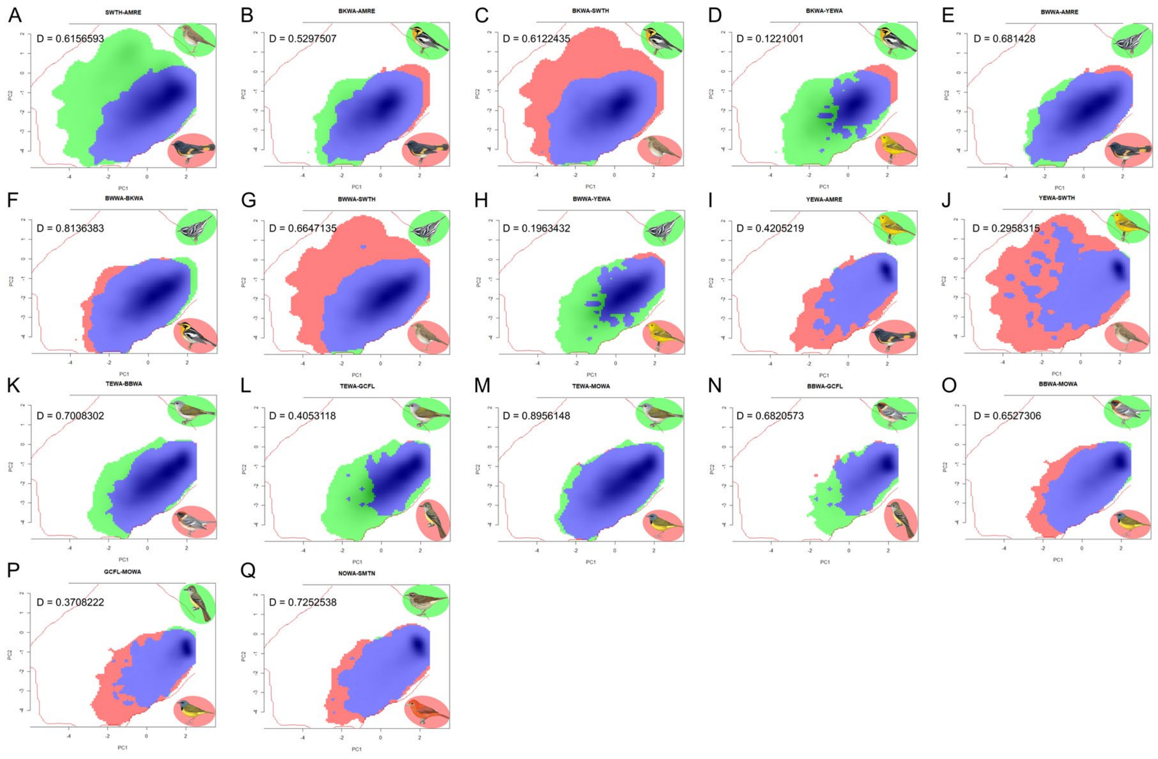


**Supplementary Figure 4.** Non-breeding climatic niche overlap between species that shared their phenological shift category. Extended Summer (A – J), Extended Winter (K – P), Late winter shift (Q). Niche overlap D values for each comparison shown in each box. Related to Figure 2 and Table S1.

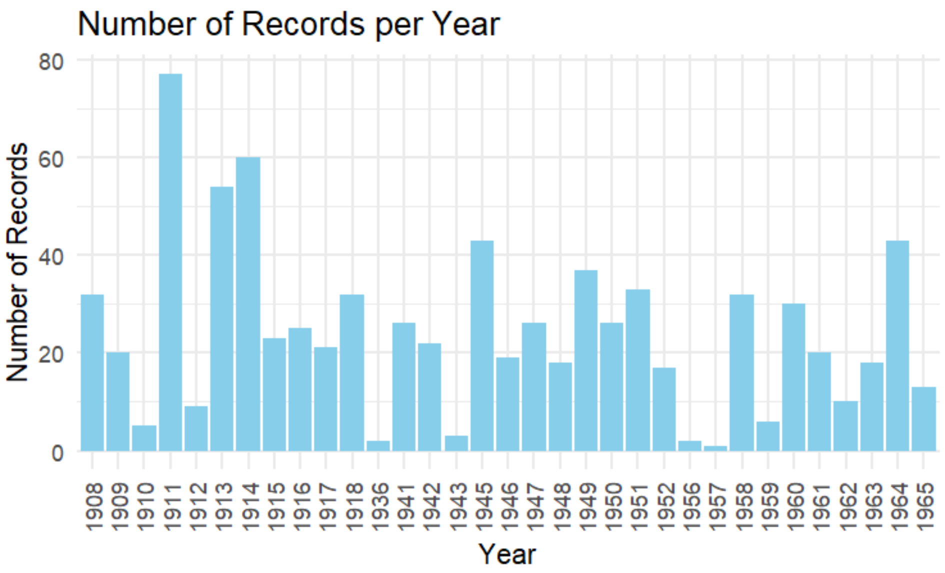


**Supplementary Figure 5.** Number of historical records available between 1908-1965, show that there is no detectable temporal bias in the availability of records. The high numbers between 1911 and 1914 correspond to the Natural History expeditions led by the American Museum of Natural History at that time (Chapman 1917). Related to STAR Methods.

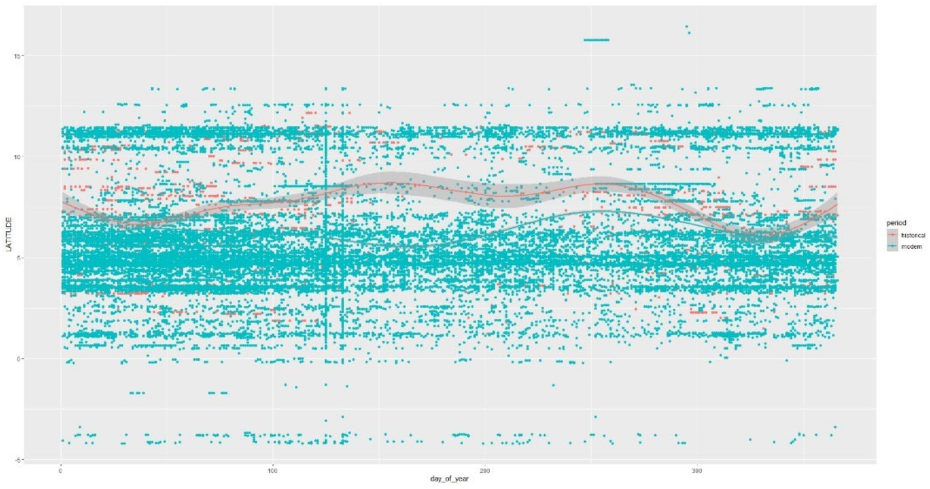

**Supplementary Figure 6.** Evaluation of the latitudinal and temporal distribution of historical (red) and modern (green) records of migratory species suggest there are no temporal and spatial biases in the data that could be influencing the patterns observed. Related to STAR Methods.

A.

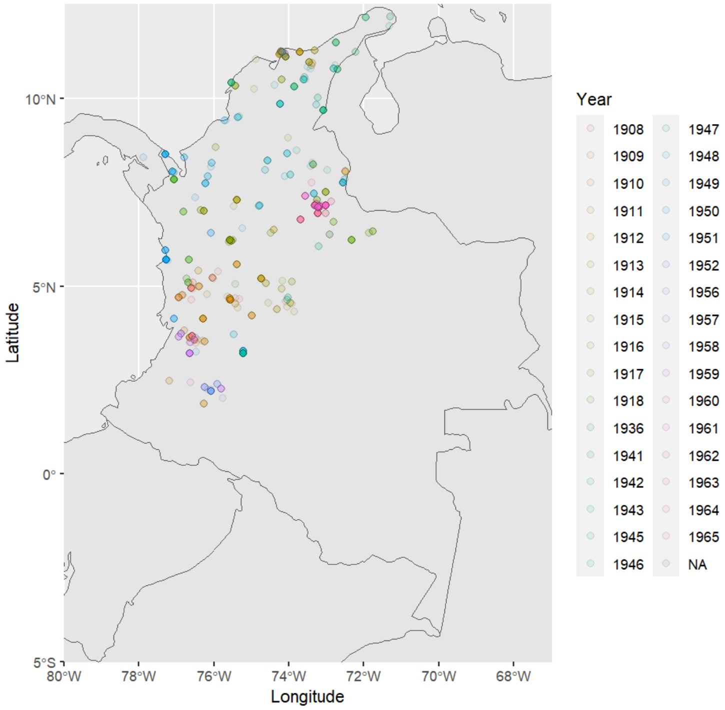


B.

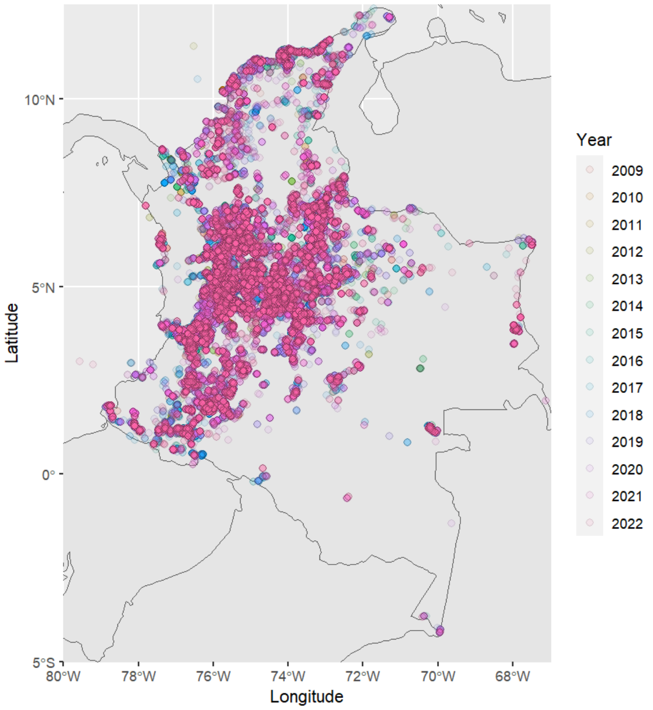


**Supplementary Figure 7.** Maps of Colombia showing the geographic distribution of migratory bird records in the A. historical dataset, and B. modern dataset. Related to STAR Methods.

**Supplementary Table 1. A.** Values of breeding climatic niche overlap ($D_{summer}$), and **B.** non-breeding climatic niche overlap ($D_{winter}$) estimated for all pairs of species in this study. Related to Figure 2 and Figure S3 and S4.

**A**

| | Group | BWWA | BKWA | YEWA | AMRE | SWTH | GCFL | MOWA | TEWA | BBWA | SMTN | NOWA | ACFL |
|---|---|---|---|---|---|---|---|---|---|---|---|---|---|
| **BWWA** | Extended Summer | 1.00 | 0.69 | 0.56 | 0.83 | 0.55 | 0.41 | 0.75 | 0.61 | 0.67 | 0.23 | 0.55 | 0.24 |
| **BKWA** | Extended Summer | | 1.00 | 0.41 | 0.68 | 0.47 | 0.27 | 0.83 | 0.50 | 0.73 | 0.07 | 0.44 | 0.09 |
| **YEWA** | Extended Summer | | | 1.00 | 0.61 | 0.67 | 0.27 | 0.48 | 0.68 | 0.44 | 0.16 | 0.76 | 0.17 |
| **AMRE** | Extended Summer | | | | 1.00 | 0.64 | 0.38 | 0.75 | 0.66 | 0.63 | 0.16 | 0.59 | 0.18 |
| **SWTH** | Extended Summer | | | | | 1.00 | 0.17 | 0.55 | 0.70 | 0.56 | 0.06 | 0.68 | 0.03 |
| **GCFL** | Extended Winter | | | | | | 1.00 | 0.27 | 0.15 | 0.17 | 0.61 | 0.16 | 0.70 |
| **MOWA** | Extended Winter | | | | | | | 1.00 | 0.60 | 0.80 | 0.05 | 0.52 | 0.05 |
| **TEWA** | Extended Winter | | | | | | | | 1.00 | 0.64 | 0.03 | 0.85 | 0.02 |
| **BBWA** | Extended Winter | | | | | | | | | 1.00 | 0.16 | 0.55 | 0.02 |
| **SMTN** | Late winter shift | | | | | | | | | | 1.00 | 0.04 | 0.82 |
| **NOWA** | Late winter shift | | | | | | | | | | | 1.00 | 0.03 |
| **ACFL** | Early winter shift | | | | | | | | | | | | 1.00 |

**B**

| | Group | BWWA | BKWA | YEWA | SWTH | AMRE | TEWA | BBWA | GCFL | MOWA | NOWA | SMTN | ACFL |
|---|---|---|---|---|---|---|---|---|---|---|---|---|---|
| **BWWA** | Extended Summer | 1.00 | 0.81 | 0.20 | 0.66 | 0.68 | 0.89 | 0.59 | 0.30 | 0.90 | 0.35 | 0.57 | 0.73 |
| **BKWA** | Extended Summer | | 1.00 | 0.12 | 0.61 | 0.53 | 0.72 | 0.44 | 0.17 | 0.74 | 0.25 | 0.45 | 0.59 |
| **YEWA** | Extended Summer | | | 1.00 | 0.30 | 0.42 | 0.33 | 0.53 | 0.74 | 0.29 | 0.81 | 0.56 | 0.39 |
| **SWTH** | Extended Summer | | | | 1.00 | 0.62 | 0.69 | 0.52 | 0.32 | 0.66 | 0.39 | 0.62 | 0.65 |
| **AMRE** | Extended Summer | | | | | 1.00 | 0.77 | 0.78 | 0.50 | 0.75 | 0.56 | 0.79 | 0.82 |
| **TEWA** | Extended Winter | | | | | | 1.00 | 0.70 | 0.41 | 0.90 | 0.45 | 0.67 | 0.82 |
| **BBWA** | Extended Winter | | | | | | | 1.00 | 0.68 | 0.65 | 0.67 | 0.78 | 0.83 |
| **GCFL** | Extended Winter | | | | | | | | 1.00 | 0.37 | 0.78 | 0.63 | 0.55 |
| **MOWA** | Extended Winter | | | | | | | | | 1.00 | 0.39 | 0.62 | 0.79 |
| **NOWA** | Late Winter Shift | | | | | | | | | | 1.00 | 0.73 | 0.57 |
| **SMTN** | Late Winter Shift | | | | | | | | | | | 1.00 | 0.79 |
| **ACFL** | Early Winter Shift | | | | | | | | | | | | 1.00 |

**Supplementary Table 2.** Values of the Durbin-Watson autocorrelation test between the number of records of migratory birds and year, longitude and latitude, both for historical and modern datasets. Values around 2 of the D-W statistic show no autocorrelation of residuals. Values closer to 0 or to 4 show a positive or negative correlation respectively and are shown in bold. Related to STAR Methods and Figures S5 – S7.

| Period | Autocorrelation | D-W |
|---|---|---|
| | Year | |
| Historical | 0.16 | 1.66 |
| Modern | 0.49 | **0.54** |
| | Latitude | |
| Historical | -0.08 | 2.15 |
| Modern | 0.10 | 1.77 |
| | Longitude | |
| Historical | -0.07 | 2.14 |
| Modern | 0.10 | 1.77 |
